# A carbohydrate-binding protein, FLOURY ENDOSPERM 6 influences the initiation of A- and B-type starch granules in wheat

**DOI:** 10.1101/643759

**Authors:** Tansy Chia, Marcella Chirico, Rob King, Ricardo Ramirez-Gonzalez, Benedetta Saccomanno, David Seung, James Simmonds, Martin Trick, Cristobal Uauy, Tamara Verhoeven, Kay Trafford

## Abstract

Previously, we identified a quantitative trait locus on the group 4 chromosomes of Aegilops and bread wheat that controls B-type starch-granule content. Here, we identify a candidate gene by fine-mapping in Aegilops and confirm its function using wheat TILLING mutants. This gene is orthologous to the *FLOURY ENDOSPERM 6* (*FLO6*) gene of rice and barley and the PTST2 gene of Arabidopsis. In Triticeae endosperm, reduction in the gene dose of functional *FLO6* alleles results in reduction, or loss, of B-granules. This is due to repression of granule initiation in late-grain development, but has no deleterious impact on the synthesis of A-granules. The complete absence of functional FLO6, however, results in reduced numbers of normal A-type and B-type granules and the production of highly-abnormal granules that vary in size and shape. This polymorphous starch seen in a wheat *flo6* triple mutant is similar to that observed in the barley mutant Franubet. Analysis of Franubet (fractured Nubet) starch suggests that the mutant A-granules are not fractured but compound, due to stimulation of granule initiation in plastids during early-grain development. Thus, in different situations in Triticeae, FLO6 either stimulates or represses granule initiation.

## Introduction

Triticeae species, such as bread wheat (*Triticum aestivum L.)*, are unusual amongst grasses in having two types of starch granule in their endosperm called A- and B-type granules. These originate from two starch granule initiation events that are separated in time and space. The first event gives rise to a single large A-type granule per plastid and takes place early in endosperm development in the main body of the plastid. The second granule initiation event gives rise to small B-type granules and takes place several days after the first event during endosperm development at least partly within the plastid stromules. Thus, endosperm plastids in wheat each contain one large A-type granule and several small B-type granules.

The endosperms of almost all species of Triticeae, including the domesticated cereals wheat, barley (*Hordeum vulgare L*.) and rye (*Secale cereale L*.), and also wild grasses such as Aegilops, contain both A- and B-type starch granules. However, a few species of wild Triticeae have normal A-granules, but lack B-granules such as *Aegilops peregrina* (Hack.) which was shown to be B-granule-less (Stoddard, 1999; Stoddard and Sarker, 2000). To understand the control of B-granule content, the impact of B-granules on starch functional properties and the mechanisms involved in starch granule initiation in wheat (and in plants generally), we created a population of Aegilops with varying B-granule content by crossing the B-less tetraploid *Ae. peregrina* with a synthetic tetraploid Aegilops called KU37 that has B-granules (Howard *et al.*, 2011). KU37 has the same genome composition (SSUU) as *Ae. peregrina* but was derived from a cross between the diploids *Ae. sharonensis* (SS) and *Ae. umbellulata* (UU) (Tanaka, 1955, 1983). Using this population, we identified a quantitative trait locus (QTL) on the short arm of chromosome 4S that accounted for 44% of the control of B-granule content (Howard *et al.*, 2011). As a result, we hypothesised that other Triticeae species, such as bread wheat, have a gene controlling <u>B</u>-<u>g</u>ranule <u>c</u>ontent, *BGC1* in a syntenous position on the group 4 chromosomes. We tested this hypothesis in the bread wheat cultivar Paragon by selecting and combining together in one plant, large chromosomal deletions spanning the putative *BGC1* regions of chromosomes 4A and 4D (Chia *et al*., 2017). The single-deletion mutants had normal starch, but the double-deletion mutant line was found to lack B-type starch granules. Thus, we successfully transferred the B-granule-less trait seen in Aegilops to bread wheat, and showed that the *BGC1* gene responsible for granule initiation must be one of the 240 genes that were common to both the 4A and 4D deletions (Chia *et al*., 2017).

Here, we describe the identification of the *BGC1* gene in Aegilops and wheat. First, we mapped *BGC1* in progeny derived from the *Ae. peregrina* x KU37 cross that had been used for the initial QTL mapping. We then tested a candidate gene using TILLING mutants of both tetraploid and hexaploid wheat (Krasileva *et al*., 2017) and confirmed its role in controling B-granule content. We found that *BGC1* is already known to be involved in starch granule initiation from work on mutants of rice (*Oryza sativa* L.), barley, and Arabidopsis (*Arabidopsis thaliana* L.). However, the phenotype of these mutants, even that of barley which is also a Triticeae, differs from the B-granule-less phenotype we have seen in Aegilops and wheat. To understand this difference, we compared the starch phenotypes of wheat lines with different dosages of functional *BGC1*.

## Results

### Fine mapping BGC1 in Aegilops

Progeny from the *Ae. peregrina* x KU37 cross were genotyped to identify plants with recombination in the *BGC1* region on chromosome 4S (Howard *et al*., 2011). Genetic markers were designed to polymorphisms between the two parents, *Ae. peregrina* and KU37, that were identified in RNA-Seq data from grains and leaves (Supplementary Table 1). Selected recombinant plants were allowed to self-fertilize and homozygous recombinant lines were identified, phenotyped and genotyped to generate a high-density genetic map across the *BGC1* interval (Fig. 1).

**Figure 1.**
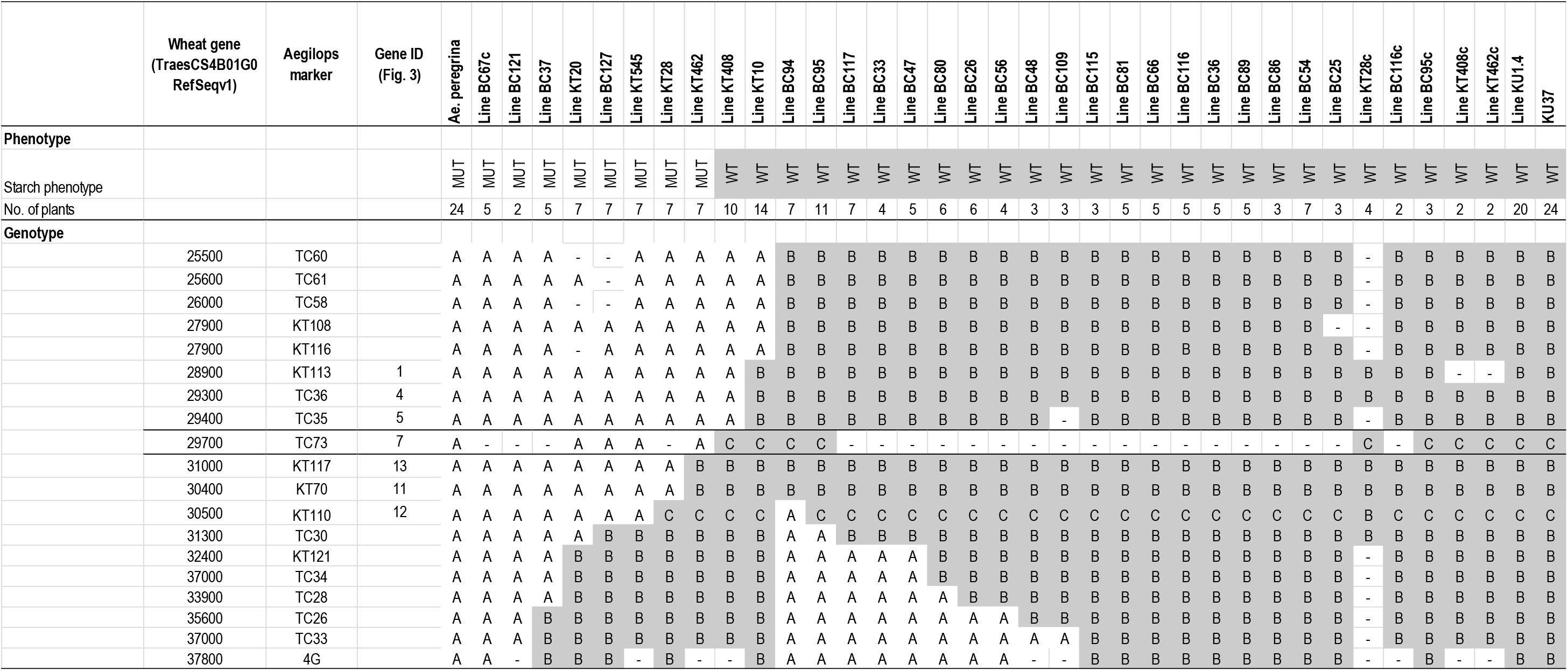
Fine-mapping B-granule content in Aegilops. Graphical genotype of the progeny derived from a cross between an Aegilops with few or no B-type starch granules, *Ae. peregrina* (mutant) and one with B-granules, KU37 (wild type). Recombinant plants were genotyped using markers within the previously identified QTL region controlling B-granule content (Howard *et al*., 2011). The grain starch was phenotyped using image analysis methods 1 or 2. The genotype codes are: ‘A’, Homozygous mutant; ‘B’, Homozygous wild type; ‘C’, either B or Heterozygous (not A); and ‘-‘, no data available. The phenotype codes are: ‘A’, lacks B-type starch granules (mutant) and ‘B’, has both A- and B-type starch granules (wild type). Line names ending in ‘c’ indicate non-recombinant sibling controls. The mapping data indicates that the gene responsible for the trait is located between the two horizontal lines in the table. Marker TC73, within this region was developed to test the candidate gene.

The homozygous recombinant lines were grown and analysed in multiple years and locations to identify lines with different granule-size distributions (Fig. 2A). The genetic and phenotypic data mapped *BGC1* between two groups of co-segregating markers, KT113/TC36/TC35 and KT117/KT70 (Fig. 1).

**Figure 2.**
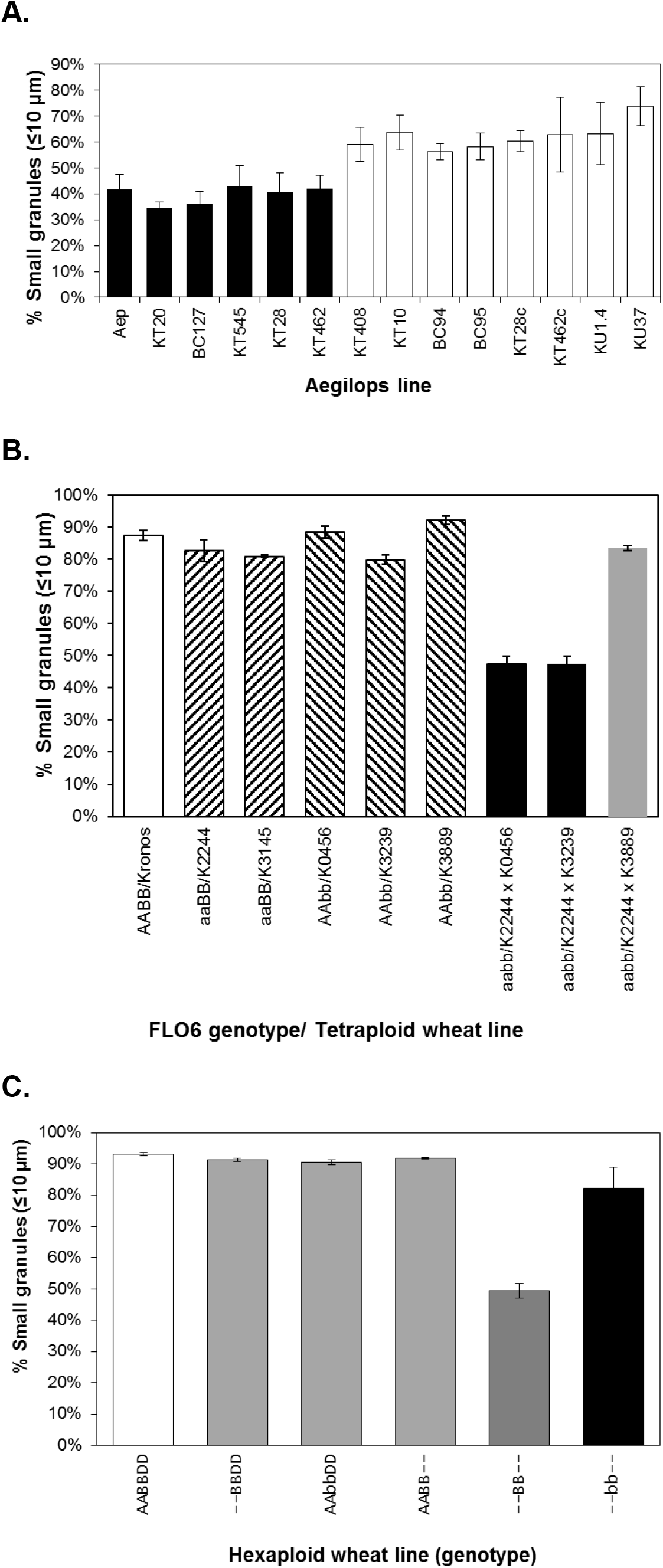
Starch granule phenotype in Aegilops and wheat. Graphs show the starch phenotype (% of granules with diameters ≤ 10 µm) of selected lines determined using image analysis method 2 (Cell Counter). A. Aegilops lines from the mapping population including the parents, *Ae. perigrina* (Aep, mutant) and KU37 (wild type). Line names ending in ‘c’ indicate non-recombinant sibling controls. Black bars = mutant genotype for marker TC73 (*FLO6-S*). White bars = wild-type genotype for marker TC73 (*FLO6-S*). Values are means ± SE for 4 to 10 samples of grain starch, each from a different plant. B. Tetraploid wheat TILLING lines. White bar = wild type Kronos. Striped bars = single mutants affected in *FLO6-A* or *FLO6-B*. Black bars = double TILLING mutants with low or no B-type starch granules. Grey bar = double TILLING mutant with wild-type starch phenotype. Values are means ± SE for 3 to 16 samples of grain starch, each from a different grain. C. Hexaploid wheat deletion and TILLING lines. White bar = wild-type Paragon. Grey bars = single mutants lines (‘--BBDD’, Paragon A-genome deletion mutant; ‘AAbbDD,’ Cadenza B-genome *FLO6-B* TILLING mutant; ‘AABB--‘, Paragon D-genome deletion mutant). Dark grey bar = Paragon double deletion mutant lacking B-type starch granules. Black bar = triple *FLO6* mutant. Values are means ± SE for 3 to 16 samples of grain starch, each from a different grain.

### FLOURY ENDOSPERM 6 is a strong candidate for BGC1

We used the *Aegilops* genetic map to establish the syntenic interval in the reference hexaploid bread wheat genome of accession Chinese Spring. Our previous analysis of bread wheat deletion mutants showed that *BGC1* is present in at least two genomes (A and D; Chia *et al*., 2017). We therefore simplified the synteny analysis by ignoring genes in the region of interest that are present in only one genome (Fig. 3). Comparison of gene order between the wheat A, B and D sub-genomes and the Aegilops S sub-genome showed only minor differences, with local re-arrangements. From this synteny analysis, we identified 13 genes as candidates for *BGC1* (Supplementary Table 2). Examination of their annotations showed that one gene, known in rice as *FLOURY ENDOSPERM 6* (*FLO6)* (Gene 7 in Fig. 3 and Supplementary Table 2), has a known involvement in starch synthesis. In rice and barley, mutations disabling the *FLO6* gene cause disruption of starch granule structure (Suh *et al*., 2004; Peng *et al*., 2014; Saito *et al*., 2018) and in Arabidopsis, disruption of the orthologous gene *PTST2* (At1g27070) causes reduction in the number of starch granules per chloroplast from 5-7 to 0-1 (Seung *et al*., 2017). This type of gene, which will hereafter be referred to as *FLO6*, encodes a protein belonging to a family of proteins in higher plants called PROTEIN TARGETING TO STARCH (PTST). All members of PTST contain two types of domain: one or more coiled-coil domains (thought to be involved in protein-protein interactions; Mason and Arndt, 2004) and a C-terminal carbohydrate binding domain of the CBM48 class (Seung *et al*., 2017).

**Figure 3.**
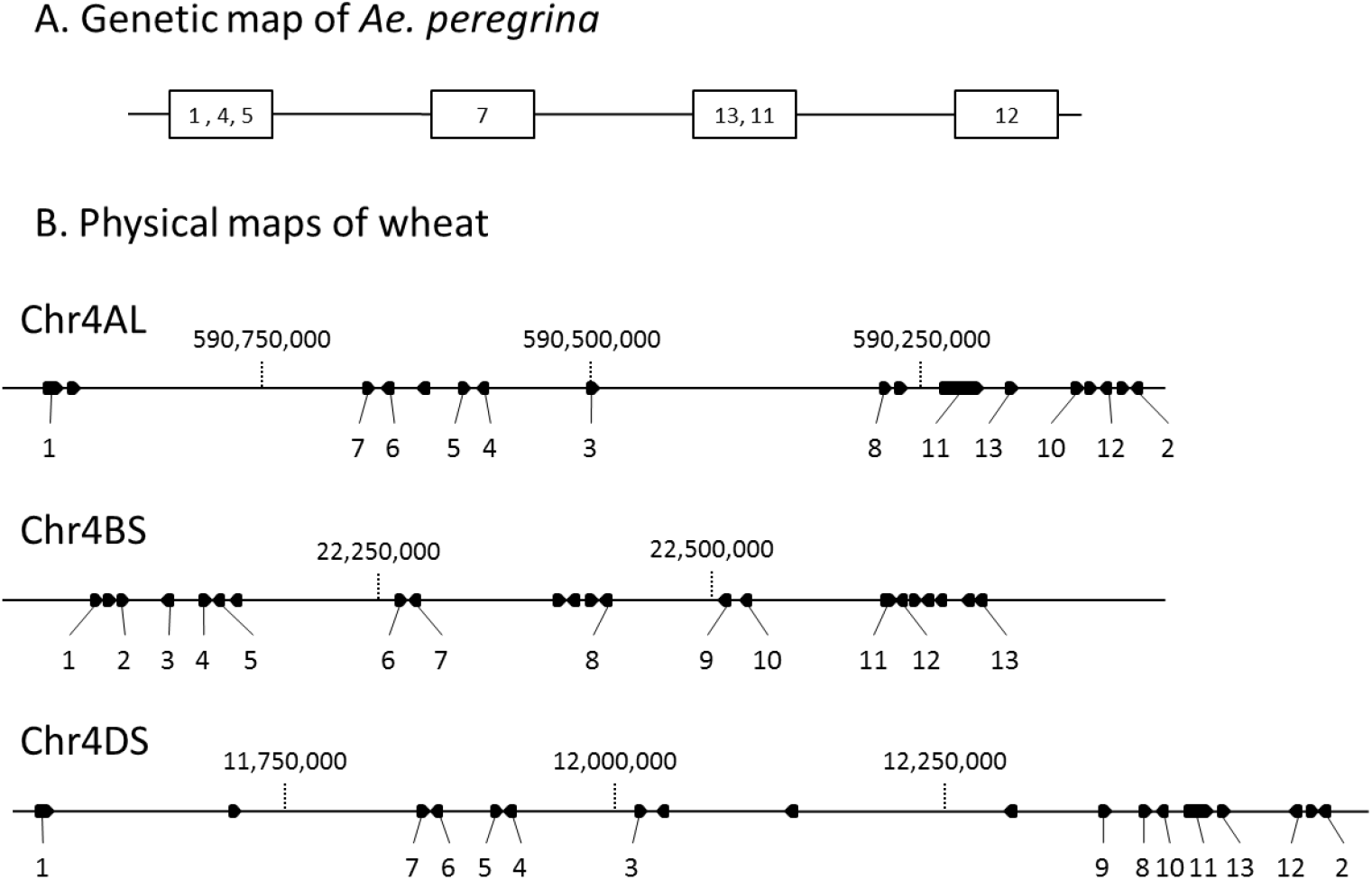
Physical and genetic maps of wheat and Aegilops. The locations of genes in the *BGC1* regions of *Ae. peregrina* and wheat are shown. Genes are numbered according to their order on chromosome 4B in wheat (see Supplementary Table 2). Orthologous genes and their corresponding markers have the same number in each map. A. Genetic order of genes in Aegilops determined from the data in Figure 1 for lines derived the *Ae. peregrina* x KU37 cross. Markers at the same genetic position are boxed. B. Physical maps across *BGC1* on chromosomes 4A, 4B and 4D of *T. aestivum* cv Chinese Spring (Refseqv1) (IWGSC, 2018). Genes and their orientations are indicated by arrows. Only genes present on all 3 chromosome-portions are numbered (below the genes). Positions in bp are indicated (above the genes).

### Sequencing the FLO6 gene in Aegilops

To test whether *FLO6* is *BGC1* in Aegilops, we cloned and sequenced the *FLO6* genes of the mutant *Ae. peregrina* and the wild-type KU37 (Supplementary Table 3A). Two homoeologous sequences representing the U and S sub-genome homoeologs of *FLO6* (*FLO6-U* and *FLO6-S*) were expected for each of these tetraploid Aegilops. Partial genomic sequences of Aegilops *FLO6* were obtained and have been deposited in GenBank (accession numbers MK848198, MK848199, MK848200, MK848201). The S- and U-genome sequences were differentiated by the SNP sequence in the *FLO6-S* gene that was used to map this gene to the *BGC1* interval (marker TC73; Fig.1), which was shown previously to lie on chromosome 4S (Howard *et al.*, 2011). Compared to rice *FLO6* (Fig. 4 and Supplementary Fig. 1), the KU37 and *Ae. peregrina FLO6-U* sequences lack the 5’ end of exon 1 that encodes the transit peptide and the first 9-13 amino acids of the mature protein. These two partial sequences vary, particularly in the intron sequences, but they encode identical proteins. We also cloned a truncated *FLO6-S* gene from KU37 that lacked exons 1 to 3 and most of the 3’ UTR. However, we were unable to clone any of the *FLO6-S* gene from *Ae. peregrina* except for the 3’ end encoding exon 8 and 9, and the 3’ UTR. This raised the possibility that in *Ae. peregrina*, the *FLO6-S* gene is almost completely deleted at the 5’ end. To explore this further, primers were designed to amplify various regions of the *FLO6* genes of Aegilops and wheat (Supplementary Table 3B and Supplementary Fig. 2A). No amplification was seen with primers binding to intron 3, exon 4-5 or intron 5 of *FLO6-S* of *Ae. peregrina* although products (Supplementary Fig. 2A; P4, P5 and P6) were obtained with these primers for KU37 *FLO6-S*. This suggests that the B-granule-less *Ae. peregrina* may contain a functional *FLO6-U* but that the other homoeolog, *FLO6-S* is probably dis-functional due to severe truncation.

**Figure 4.**
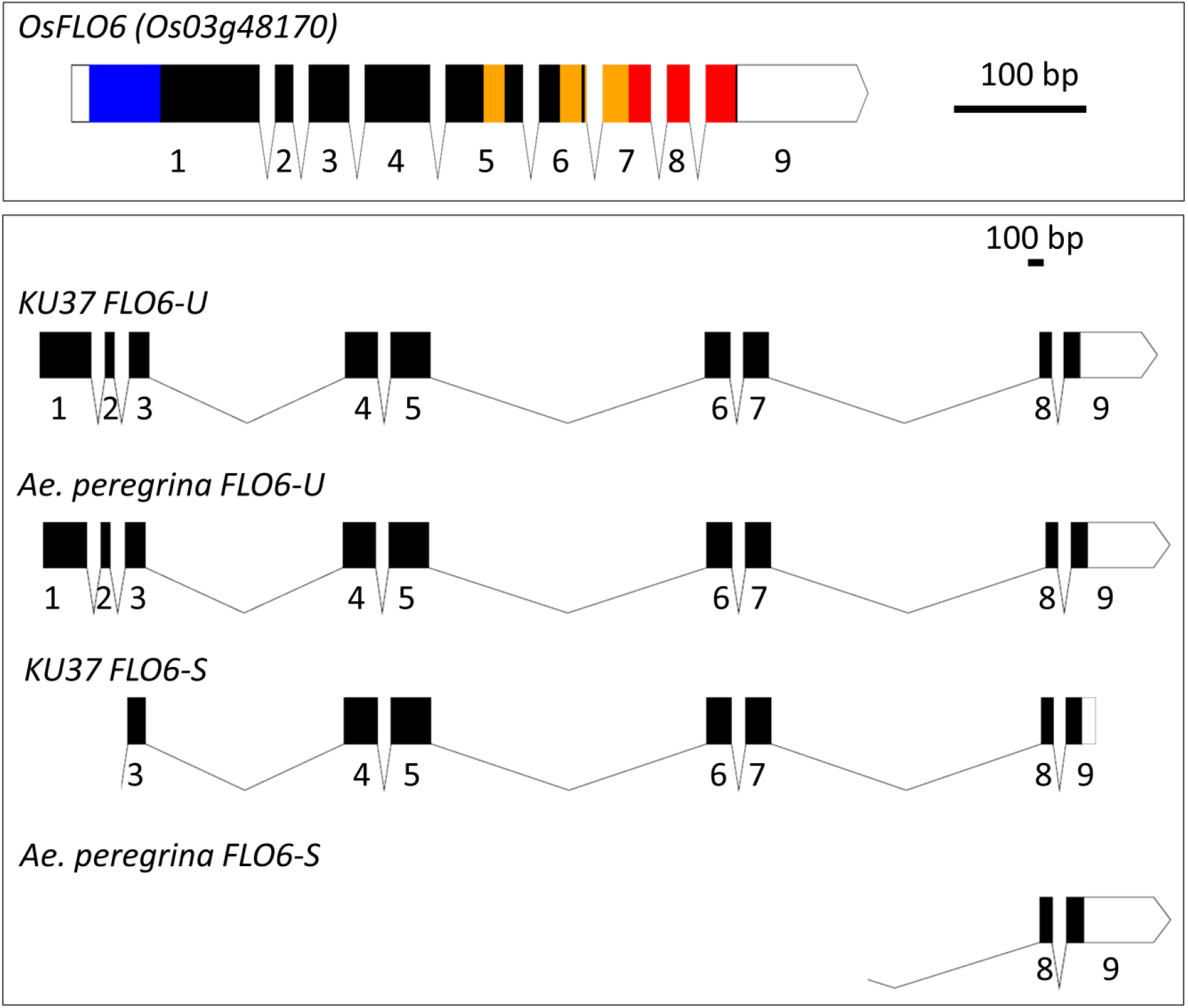
*FLO6* gene structure in rice and Aegilops. Diagrams of the gene structures are drawn to scale (bar = 100bp). The exons are shown as filled boxes and numbered 1 to 9. The 5’- and 3’-UTRs are shown as unfilled boxes. For rice FLO6, the sequence encoding the transit peptide is coloured blue, the coiled coil domains yellow and the starch binding domain red.

### Testing the candidate gene, FLO6 in wheat

To test whether mutations in the wheat *FLO6* gene result in reduction or elimination of B-type starch granules, we selected TILLING lines of the *T. durum* wheat cultivar, Kronos (a tetraploid with genome composition AABB) with EMS-induced mutations in either the A-genome (*FLO6-A*) or B-genome (*FLO6*-*B*) homoeolog (Krasileva *et al.*, 2017; www.wheat-tilling.com; Supplementary Figure 3). Both of the *FLO6-A* lines (Kronos2244 and Kronos3145) have induced nonsense mutations resulting in premature stop codons. All of the *FLO6*-*B* lines (Kronos3239, Kronos0456 and Kronos3889) have missense mutations in the region encoding the FLO6 CBM48 domain.

Starch from homozygous single-mutant grains was subjected to image analysis to quantify the small-granule content (B-granules plus small A-granules) (Fig. 2B). All of the single mutant lines had a normal small-granule content (Student’s t-test; P > 0.05). One of the *FLO6*-*A* nonsense lines, K2244 was crossed to each of the three *FLO6*-*B* missense lines and homozygous double mutant lines (and their corresponding wild-type segregant lines) were selected. One of the double mutants (K2244 × K3889) had a normal small-granule content, but the other two (K2244 × K3239 and K2244 × K0456) had significantly lower (54% to 59%) small-granule contents than the wild types and the single mutants. When observed microscopically, the grains and extracted starch from these two double mutants had very few B-granules (e.g. Fig. 5A; *T. durum* aabb; K2244 × K3239). The small-granule content of these two tetraploid (Kronos) wheat double mutants (Fig. 2B) was very similar to that seen for the hexaploid (Paragon) double-deletion mutant (--BB--; Fig. 2C) and the natural B-granule-less species *Ae. peregrina* (Aep; Fig. 2A) that were described previously (Howard *et al.*, 2011; Chia *et al.*, 2017).

**Figure 5.**
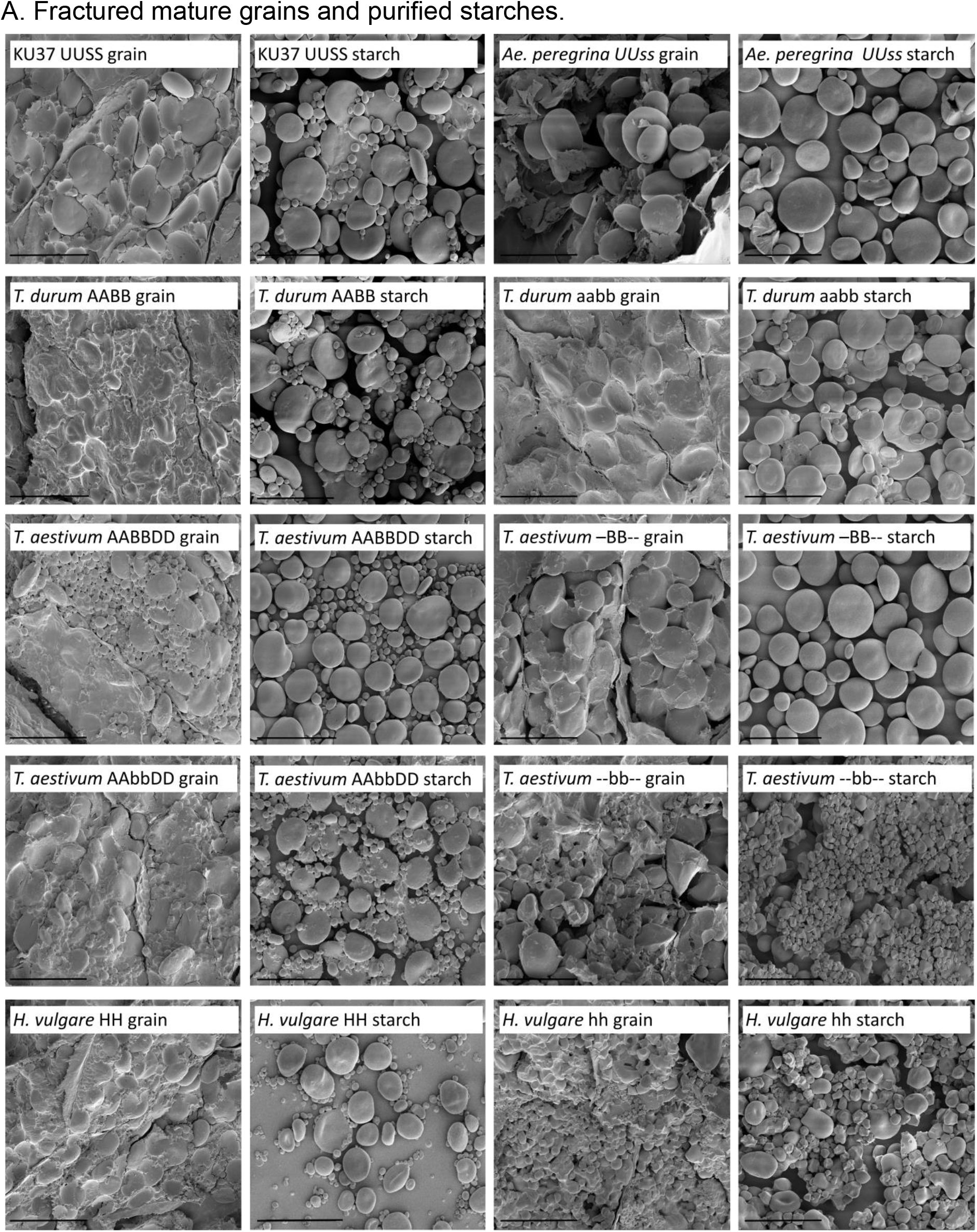
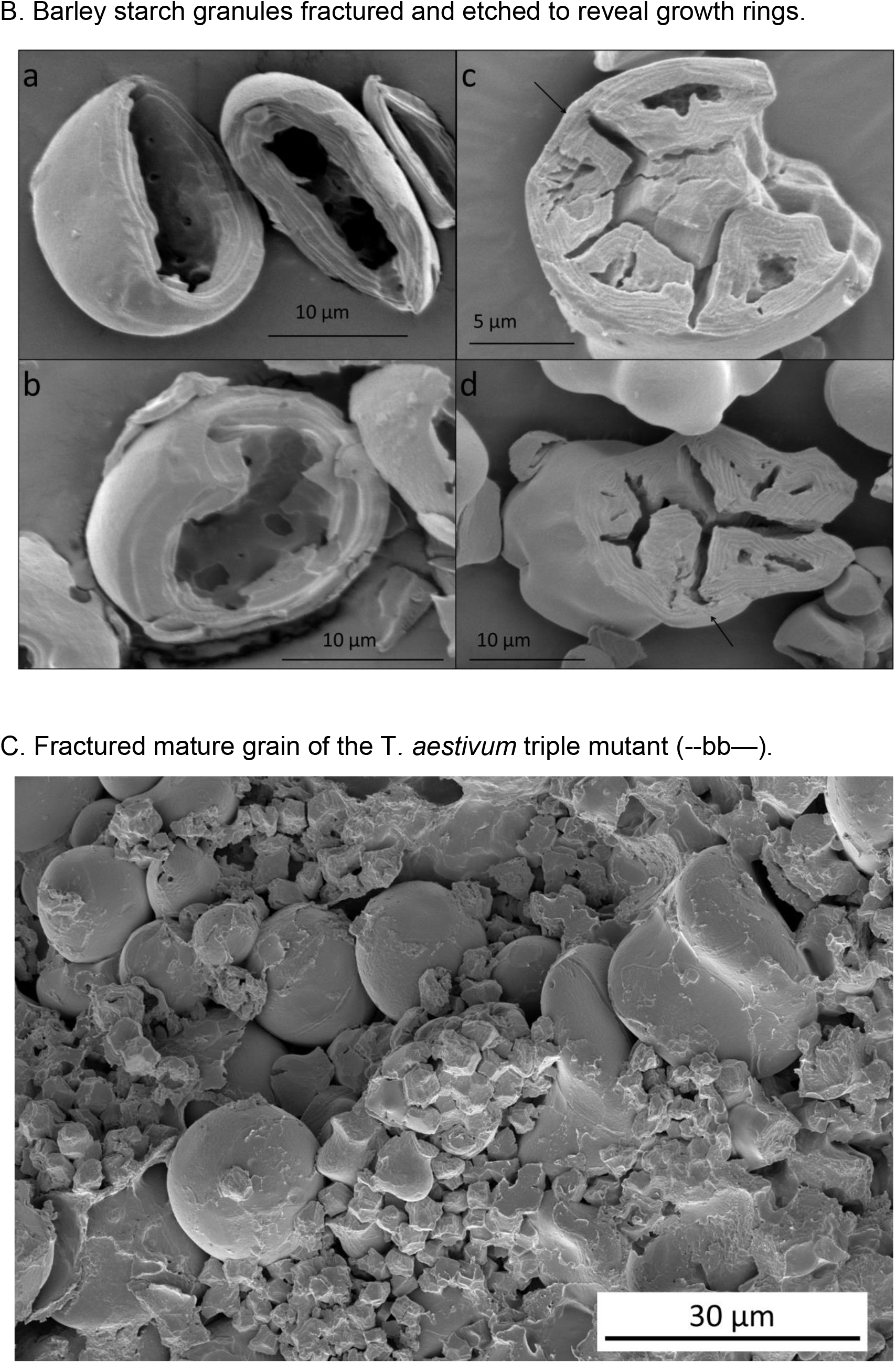
Scanning Electron Microscope images of starch and grains. A. Fractured mature grains and purified starches For each image, the species, the genome composition, the genotype of FLO6 (Upper case genome letter for wild type, lower case for mutant, - for deletion) and the source of the sample (grain or starch) is shown. Scale bars are 50 µm. B. Barley starch granules fractured and etched to reveal growth rings. The growth-ring structure within the starch granules was revealed by cracking starch by grinding and then partially digesting with amylase. The granules in wild-type Nubet starch (a, b) each consists of a single ring structure suggesting a single initiation point. Some of the starch granules in the mutant Franubet are compound and contain multiple separately-initiated sub-granules, each with their own ring structure. In some granules (c, d), an outer layer of starch with continuous growth rings can be seen surrounding the compound granule (indicated by the arrows) suggesting that Franubet also contains some semi-compound granules. C. Fractured mature grain of the *T. aestivum* triple mutant (--bb--) An example of the polymorphous starch observed in the triple mutant endosperm. Large, sometimes distorted, simple granules were observed as well as compound (angular) starch granules. Some cells close to the aleurone contained A-granules but no B-granules (not shown). Bars indicate the magnification.

### The barley FLO6 mutant, Franubet forms compound starch granules

Despite being closely related taxonomically, the starch phenotype of the barley *FLO6* mutant, Franubet is very different from that of the other Triticeae *FLO6* mutants (tetraploid wheat, hexaploid wheat *and Ae. peregrina*), all of which have low or zero B-granule contents. In Franubet barley, both A and B-type starch granules are largely absent and in their place are starch granules that appear to be fractured or fragmented (De Haas, Goering and Eslick, 1983).

To investigate this apparent discrepancy, we examined the grains and starch of the Franubet mutant of barley (Fig. 5A; *H. vulgare hh*). Franubet starch was clearly distinct from that of B-granule-less wheat. The starch granules were highly heterogeneous in morphology, as described previously (Verhoeven, 2002, De Haas, Goering and Eslick, 1983; Saito *et al.*, 2018). There were few, if any, normal A-type or B-type granules in Franubet starch. Some abnormally-large A-type granules, irregular or lobed granules and granules that appear to be compound or fractured were seen.

Compound granules form from multiple, separately-initiated granules that become compressed together within a plastid to form polygonal shapes. The presence of compound granules in Franubet in place of single A-type granules would indicate an increase in granule number per plastid rather than a decrease as seen in B-granule-less wheat. We therefore investigated the structure of these starch granules further. Each sub-granule in a compound granule has its own set of growth rings (unlike fractured granules that form when a single granule breaks apart late in grain development). Cracking and partly-digesting extracted Franubet and Nubet (the wild-type parent of Franubet) starch confirmed that Franubet contains compound starch granules with individual ring structures (Fig. 5B). Furthermore, examination of some of the larger-than-normal, A-type granules in Franubet showed that, although they appear simple from the outside, when cracked and etched, these granules contain within them separately-initiated sub-granules. Such granules are called semi-compound granules and have been observed in other plant species e.g. the bulbs of *Scilla ovatifolia* Baker (family Hyacinthaceae) (Badenhuizen, 1965). However, as far as we are aware, semi-compound granules have not been observed previously in cereal endosperm.

### Generation and analysis of wheat entirely devoid of functional FLO6

The apparent discrepancy between the phenotypes of wheat/Aegilops and barley *FLO6* mutants could be due to incomplete elimination of functional FLO6 protein in the former species. Unlike barley, which is diploid, the wheat and Aegilops B-granule-less mutants are polyploid. We hypothesised that not all of the *FLO6* homoeologs in these polyploid mutants are completely defective. To investigate this, we crossed the Paragon double deletion mutant (*FLO6* genotype --BB--) to a hexaploid wheat (Cadenza) TILLING mutant (Cadenza1730) with a nonsense mutation in the B-genome homoeolog of *FLO6* (genotype AAbbDD). We selected from the F_2_ progeny a triple mutant line (genotype --bb--) unable to make any FLO6 protein due to deletion of *FLO6 4A* and *FLO6 4D*, and a nonsense (STOP) mutation in *FLO6 4B*.

The starch of this *flo6* triple mutant (--bb--), when viewed microscopically, looked very different from that of the B-granule-less *Ae. peregrina*, the Paragon double deletion mutant (--BB--) and the Kronos double mutants (aabb). Rather than B-granule-less, the starch resembled that in Franubet barley (Fig. 5A; *T. aestivum*; --bb--; and Fig. 5C). However, there were more simple granules in the triple mutant wheat than in Franubet barley. These could be A-type granules or semi-compound granules. No, or very few, B-type granules were observed. The starch granules in the wheat *flo6* triple mutant were very heterogeneous in shape and size. Scanning electron microscopy (SEM) of grains showed that the granules size and shape varied greatly between cells, and between plastids with in the same cell. There were many compound granules, but these varied in the number of constituent sub-granules. Some had just a few large sub-granules, sometimes arranged in a line or in an irregular shape (these were also observed in Franubet starch). Some of the compound granules were composed of many polygonal sub-granules, similar in size to B-granules and some had many, even smaller sub-granules.

### Coulter counter analysis of starch-granule-size distribution

To further characterise the size distribution of the various wheat starches, we used a Coulter counter. We compared starches from wild-type Paragon (*FLO6* genotype AABBDD), B-granule-less Paragon double-deletion mutant (--BB--) and the triple mutant with polymorphous starch (--bb--). The starches in all three genotypes had different starch granule size distributions (Fig. 6A, B). The wild-type starch had a bimodal size distribution (Fig. 6A). By fitting a mixed gaussian distribution to the data, we deconvoluted the two overlapping peaks of A- and B-type granules, which showed that the B-granule content of Paragon starch was 4.2% ± 0.8% (by volume). The double deletion mutant starch had a unimodal size distribution as it entirely lacked the B-granule peak. The polymorphous starch in the triple mutant also had a unimodal distribution but this was wider than that of the B-granule-less starch, and included many small granules of less than 5 µm in diameter (Fig. 6B).

**Figure 6.**
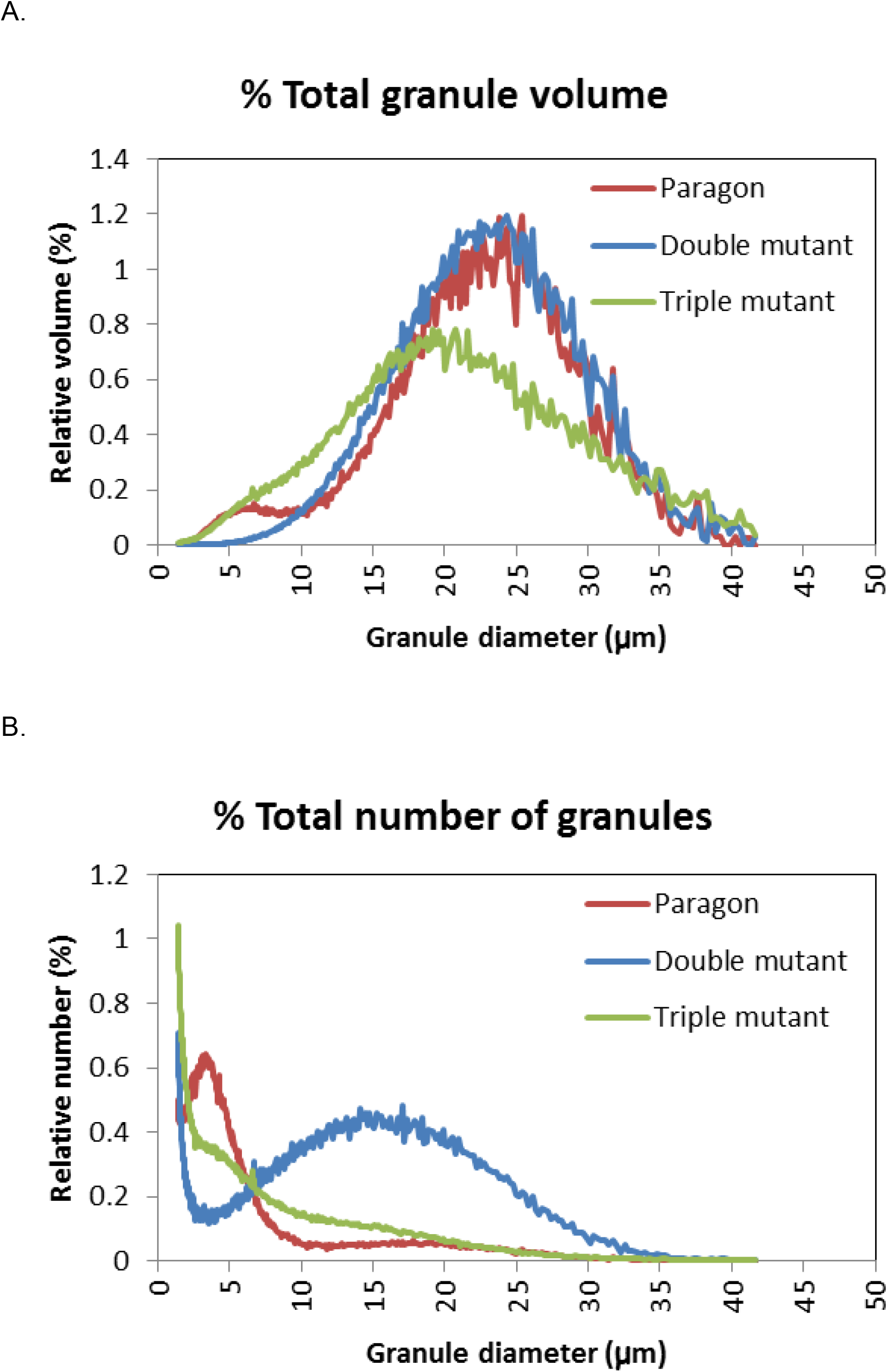
Starch granule-size distribution in hexaploid wheat. Starches were purified from mature grains of Paragon (AABBDD, wild type control), The Paragon double deletion mutant (--BB--) and the triple mutant selected from the progeny of a cross between the Paragon double deletion mutant with a Cadenza TILLING mutant with a nonsense mutation in FLO6-B (AAbbDD). Starches were analysed using a coulter counter. Values for Paragon and the Paragon double mutant are means for 3 samples of starch, each from a different grain. Values for the triple mutant are means for 6 samples of starch each from a different F_2_ grain. Data are presented in two ways: as % total granule volume (A) and as % total number of granules (B).

## Discussion

Fine mapping in Aegilops revealed a candidate gene underlying the *BGC1* locus controlling B-type starch-granule content. The candidate gene is an ortholog of the *FLOURY ENDOSPERM 6* (*FLO6*) gene in rice (Peng *et al*., 2014). Cloning and sequencing of the *FLO6* genes in Aegilops supported the idea that *BGC1* is an ortholog of *FLO6*. Whilst a near-full length *FLO6-S* gene that would encode a protein with no obvious defects was found in KU37 which has normal starch, we were unable to amplify a full-length *FLO6-S* from the natural B-granule-less species *Ae. peregrina*. We suggest that the *FLO6-S* gene in *Ae. peregrina* may have been partially deleted, or substantially modified from normal. The candidate gene was further tested by selecting a tetraploid wheat mutant with disrupted *FLO6* genes. The mutant was found to have few, if any B-granules, like *Ae. peregrina* and a hexaploid bread wheat (Paragon) double-mutant with deletions of the *BGC1*-regions of chromosomes 4A and 4D (Chia *et al*., 2017). This result confirms that *BGC1* is an ortholog of *FLO6* and shows that *FLO6* controls B-granule content in wheat and Aegilops.

All of the B-granule-less wheat and Aegilops described here are polyploids and all are, or could be, partial *FLO6* mutants. Firstly, we found that *Ae. peregrina* has one *FLO6* homoeolog that is probably truncated and therefore dysfunctional and a second apparently normal *FLO6* homoeolog. Secondly, the B-granule-less hexaploid wheat double mutant lacks *FLO6*-A and *FLO6*-D but has *FLO6-B*. Thirdly, the two B-granule-less tetraploid wheat double mutants each have a *FLO6-A* nonsense mutation and a *FLO6-B* missense mutation. We cannot be sure that the two different missense mutations used here eliminate *FLO6* functionality: they may just reduce it. One other B-genome *FLO6* missense mutation (K3889) when combined with the same A-genome *FLO6* nonsense mutation had no impact on starch granule phenotype. We assume therefore that the K3889 *FLO6* missense mutation has little or no impact on FLO6 functionality.

The starch phenotypes of the *FLO6* mutants of rice and barley, which are diploid species, differ from those of Aegilops and wheat. Rice and barley *FLO6* mutants have very abnormal granules of diverse size and shape. We have referred to this type of starch as ‘polymorphous’ starch. Unlike the B-granule-less *FLO6* mutants, these diploids are thought to lack *FLO6* functionality entirely. We tested the effect of complete absence of *FLO6* in polyploid wheat by selecting a *FLO6* triple mutant of hexaploid wheat. This mutant contained deletions of *FLO6-A* and *FLO6-D* and a nonsense (stop) mutation of *FLO6-B* and is therefore unable to produce any normal FLO6 protein. The triple *FLO6* wheat mutant had polymorphous starch very similar to that seen in Franubet barley (Fig. 5) and *flo6* rice. Its starch phenotype (with abnormal granules of diverse size and shape, including compound granules) was distinctly different from that of the Paragon double-deletion mutant (which was B-granule-less).

Together these results suggest that in Triticeae species including wheat, 1) small reductions in FLO6 protein have little or no impact on granule size or shape, since single-mutants of tetraploid and hexaploid wheat have normal, or near normal, starch, 2) significant reduction in FLO6 protein results in lack of B-type starch granules but does not affect A-type granules, and 3) complete elimination of FLO6 protein results in the lack of B-type starch granules and severe disruption of A-type granule number and morphology. The precise relationship between *FLO6* gene-dose and phenotype may differ from species to species. In polyploid species, it will depend on the relative contributions of the different sub-genome homoeologs and the functionality of the proteins they encode. In missense mutants, it will depend on the extent of the deleterious effect of the mutation on FLO6 protein functionality. Consistent with this, barley grains heterozygous for the *FLO6* mutation have normal starch granule morphology (Saito *et al.*, 2018) suggesting that a reduction in FLO6 to less than 50% of the normal amount is required to disrupt starch granule morphology.

The effect of *flo6* mutants on starch granule size and number in cereal endosperm suggests that this protein impacts on the number of granule initiations per plastid. However, *flo6* mutants can have either increased or decreased numbers of initiations depending on the species and granule morphology can also vary between neighbouring endosperm cells and even between plastids in the same cell. In rice, which normally has compound granules, *flo6* mutants have at least some compound granules with more and smaller sub-granules than normal, suggesting that in wild-type rice endosperm, FLO6 can restrict the number of granule initiations per plastid. In the Triticeae, *flo6* mutants entirely lacking FLO6 have compound and semi-compound granules instead of the single A-type granule per plastid that normally forms early in grain development. Thus, one of the functions of FLO6 in wheat grains may be to limit the number of granule initiations that occur in a plastid so that a single A-type granule can form in the main body of the plastid. However, reduction but not elimination of FLO6 in the Triticeae polyploids results in the loss of B-type starch implying that FLO6 stimulates B-granule initiation in the stromules. Thus, early in grain development in wild-type Triticeae, FLO6 restricts granule initiation to one A-type granule per plastid in but then later in development, it has the opposite effect: FLO6 promotes the initiation of many B-type granules per plastid.

At present we cannot explain the apparently contradictory effects of FLO6 on granule initiation at different developmental stages, or the great diversity of starch granule morphology that is observed in the absence of FLO6. The answers may lie in understanding the timing of expression of FLO6 during grain development, its location within plastids and the nature of its interactions with starch, glucans and other proteins. Several interaction partners have been identified in Arabidopsis leaves for the FLO6 ortholog PTST2, including Starch Synthase 4, which is also required for proper starch granule initiation (Roldan *et al.*, 2007; Seung *et al*., 2017). PTST2 also co-purified in immunoprecipitation experiments with two other plastidial coiled coil proteins: MRC/PII1 and MFP1 (Seung *et al.*, 2018). MRC/PII1 is a direct interaction partner of SS4, while MFP1 is a thylakoid-associated protein that localises PTST2 to discrete patches in the chloroplast (Seung *et al*., 2018; Vandromme *et al*., 2019). The location of PTST2 in these patches was proposed to restrict granule initiation events to defined areas of the plastid. Wheat has orthologs of all of these proteins, but their role in the endosperm has not been studied. Interestingly, all these proteins appear to promote granule initiation in Arabidopsis leaves, rather than repress it. Future work may determine whether any of these interactions or localisation patterns are conserved in amyloplasts of the Triticeae, and other cereals.

We conclude from this work that FLO6 participates in controlling B-type starch granule initiation in Triticeae endosperm but that its precise effect on granule size and number varies with gene dose and stage of development. It is likely that the production of B-granule-less starch involves a delicate balance between the amount of FLO6 and that of one or more of its interacting partners. This may be easier to achieve in a polyploid species than in a diploid.

## Methods

### Plant material

The sources of the Aegilops and wheat deletion mutants, and methods for growth in glasshouse and field were as described previously (Howard *et al.*, 2011; and Chia *et al.*, 2017) except that Aegilops grown in the glasshouse were not vernalized. For fine mapping, in addition to the F_2_- and F_3_-derived (KT) lines from the cross between *Ae. peregrina* and the synthetic Aegilops, KU37 (Howard *et al.*, 2011), a set of ^backcrossed (BC) lines were developed from a cross between one F_2_ line, KU1.4,^ which has B-type starch granules and the natural B-granule-less parent, *Ae. peregrina*. One BC_1_F_1_ plant was backcrossed to *Ae. peregrina* and the BC_2_F_1_ grains were grown and screened with markers flanking the *BGC1* region (between the QTL peak marker, 4G and the telomere) (Supplementary Table 1) to identify ten heterozygous recombinant plants. These were grown and self-pollinated to give BC_2_F_2_ grains which were screened again as before; heterozygous recombinant lines were self-pollinated and homozygous recombinant lines were selected.

TILLING mutants with polymorphisms in the genes of interest in the tetraploid wheat Kronos and the hexaploid wheat Cadenza (Krasileva *et al.*, 2017) were identified from http://www.wheat-tilling.com/ and http://plants.ensembl.org/Triticum_aestivum/ and obtained from www.seedstor.ac.uk. The mutations were chosen based on their likelihood to affect the function of the encoded protein i.e. they were nonsense (stop) or missense mutations, or mutations that affect splicing sites of the mRNA (to remove introns).

Kronos and Cadenza TILLING mutants were grown in speed-breeding conditions (21-22°C, 22 h day/17-18°C, 2 h night; Watson *et al.*, 2018) in Levington’s M2 compost (LBS, Colne, Lancashire) in either a glasshouse where natural lighting was supplemented by high-pressure sodium lamps or an Adaptis 1000 growth chamber with a red/white LED canopy (Conviron, Winnipeg, Canada). Spikes were harvested four weeks post anthesis, dried at 30°C in an oven prior to sowing.

### DNA and RNA preparation

DNA was extracted from seedling leaves as described by Fulton, Chunwongse and Tanksley (1995). RNA was extracted from flag leaves and developing grains both harvested 19-21 days after ear emergence (mid grain development). RNA was extracted using an RNeasy Plant mini kit (Qiagen.com) according to the manufacturer’s instructions. Samples of RNA from *Ae. peregrina* grain, *Ae. peregrina* leaves, KU37 grains and KU37 leaves were pooled to give four samples with a total of 5 µg of total RNA per sample (at a minimum concentration of 25 ng/µl). These were submitted for sequencing to The Genome Analysis Centre (now the Earlham Institute), Norwich. After sample quality control, Illumina barcoded RNA TruSeq libraries were constructed and the four libraries were sequenced over two lanes, in pools of 2, on the Illumina HiSeq 2000 platform. Using 100-bp paired-end reads, at least 100 million reads per lane were generated. After data quality control, base calling and formatting, the Aegilops sequence data were aligned to a *T. aesivum* reference sequence (either pseudo-chromosomes that were organized using wheat-Brachypodium synteny or the NimbleGen wheat exome capture probe set). When it became available, the sequence data were re-aligned to Refseq v1.0 (IWGSC, 2018) with HISAT-2.0.5 (Kim *et al.*, 2015). The Refseq v1.0 alignments where sorted and the candidate regions where extracted (chr4A:589084002-591920577, chr4B:20715580-23835481,chr4D:10926756-13253764) using samtools-1.4.1 (Li *et al*., 2009). All RNAseq data were visualised and compared using IGV software (Robinson *et al*., 2011) to identify SNPs suitable for the design of molecular markers for use in genetic mapping. All raw RNA-Seq reads have been deposited in the Sequence Read Archive (SRA) under accession ERS3409626.

### Aegilops genotype analysis

To distinguish the parental and heterozygous plants within the mapping population, KASP primers were designed to polymorphisms in genes within the region of interest. These were tested on a sub-set of the population to see whether or not they amplified genes linked to *BGC1* using the PCR-based KASP genotyping assay as described by the manufacturer (https://www.lgcgroup.com). As the Aegilops plants used here are tetraploid and *BGC1* lies on sub-genome S but not on U, only half of the successful KASP assays would be expected to be linked to *BGC1*, and this was found to be the case (data not shown). The *BGC1*-linked markers on chromosome 4S and their corresponding genes in wheat are given in Supplementary Table 1.

### Phenotype analysis

Starch granules were extracted from single or half-grains as described previously (Chia *et al.*, 2017) except that the method was scaled down from 10 grains per sample to one half-grain per sample. Starch granule-size distribution was measured using either 1) a manual image-analysis method described previously (Chia *et al*., 2017); 2) an automated image analysis method using a cell counter (Biorad TC20) or 3) using a Coulter counter (Beckman Coulter Multisizer 4e) fitted with a 70 µm aperture tube, with Isoton II diluent (Beckman Coulter). Methods 1 and 2 both involve microscopic examination of starch followed by image analysis to define the proportion of granules <10 µm in diameter. B-type starch granules in Triticeae species are generally found to be smaller than 10 µm in diameter whereas A-type granules are 5-40 µm in diameter (Fig. 5). Thus, the sizes of A- and B-type granules overlap and the <10 µm category contains both B-granules and some small A-granules. For clarification, these measurements will therefore be referred to as measurements of small granules rather than measurements of B-granules.

Method 3, the Coulter counter, estimates particle volume from the transient drop in electrical resistance caused by the passage of a particle through an aperture. For samples containing both A- and B-type granules, the relative total volume of B-type starch granules was estimated by fitting a mixed gaussian distribution. A minimum of 50,000 particles were measured per sample.

### Cloning and sequencing Aegilops FLO6

Primers were designed to FLO6 genes from *Ae. tauschii* or *T. aestivum* and used to amplify the Aegilops FLO6 genes in overlapping fragments (Supplementary Table 3A). Suitable fragments were gel purified and cloned into *E.coli*, and individual clones isolated and multiplied using a CloneJet PCR cloning kit and GeneJet plasmid kit (both from Thermofisher.com), according to the manufacturer’s instructions. Multiple (>3) clones for each gene were sequenced and the trimmed consensus sequences were assembled into contigs using Geneious software (www.geneious.com). The intron/exon positions were inferred by comparison with the rice *FLO6* cDNA sequence (LOC_Os03g48170).

We were unable to amplify fragments of FLO6-S spanning exons 1-7 from B-granule-less *Ae. peregrina* using our cloning primers. To test this region further, several more 4S-specific primer pairs (and various controls) were designed and used to amplify genomic DNA from Aegilops and wheat (Chinese Spring). The primer sequences, PCR conditions and associated amplicon information are given in Supplementary Table 3B and Supplementary Figure 2.

### Genotype analysis of TILLING mutants of wheat

Primers (Supplementary Figure 3) were designed to distinguish mutant and wild type sequences and the primary plants and their progeny were genotyped using the PCR-based KASP™ genotyping assay as described by the manufacturer (https://www.lgcgroup.com). Phenotyping was performed as described above for Aegilops grains (method 2).

### Identification of triple FLO6 mutants in hexaploid wheat

The Paragon double-deletion mutant (with deletions of the FLO6 regions of chromosome 4A and 4D; Chia *et al.*, 2017) was crossed to a Cadenza TILLING mutant Cadenza1730 with a nonsense mutation in FLO6-B (Q184*). The F1 grains were allowed to self-pollinate, the resulting F_2_ grains were cut in half and the embryo-halves were grown. DNA was extracted from the F_2_ seedling leaves and this was genotyped to identify triple-mutant F_2_ plants as follows. Plants homozygous for the TILLING mutation in *FLO6-B* were identified using a KASP genotyping assay (Supplementary Table 4). Plants homozygous for either the A-genome or the D-genome deletions were identified by the absence of PCR amplification with homoeolog-specific primers to *FLO6-A* and *FLO6-D*. As a positive control, DNAs from putative deletion mutants were also tested by PCR with primers specific for *FLO6-B* and unaffected by the TILLING mutation. Starch was extracted from the F_2_ half-grains identified as triple mutants, and examined by microscopy.

### Scanning electron microscopy

Samples of purified starch were suspended in water and 2 µl was applied to a glass cover slip mounted on the surface of an aluminium pin stub using double-sided adhesive carbon discs (Agar Scientific Ltd, Stansted, Essex, UK). The water was allowed to evaporate. Mature grains were fractured in mid-section using a scalpel blade and mounted directly on double-sided adhesive carbon discs attached to stubs. The stubs were sputter coated with ~15 nm gold particles in a high-resolution sputter-coater (Agar Scientific Ltd), transferred to a FEI Nova NanoSEM 450 and viewed at 3 kV.

Fractured and etched samples of Franubet barley starch were prepared for SEM as follows. Extracted starch (0.4 g) was suspended in 1 ml dH_2_O, frozen in liquid nitrogen and ground with a pestle and mortar pre-cooled with liquid nitrogen until the starch slurry began to thaw. The mortar was refilled with liquid nitrogen and the starch slurry was ground again. This procedure was repeated once more. The ground starch slurry was transferred to1.5-ml tubes, centrifuged for 2 min at 5700 *g* and the supernatant was discarded. The pooled pellets were resuspended in a total of 3 ml dH_2_O and aliquots of 0.5 ml were added to 0.5 ml 100 mM sodium acetate (pH 5.5) containing 0 (control) or 89 U α-amylase (sigmaaldrich.com). After incubation at 37 °C for 0 (control) or 2 to 3 h and centrifugation at 5700 g for 2 min, the supernatant was discarded and 0.5 ml cold (−20 °C) acetone was added. The samples were allowed to settle for 12 h and to air-dry prior to SEM preparation and observation as above.

## Acknowledgements

We thank the International Wheat Genome Sequencing Consortium (IWGSC) for pre-publication access to the RefSeq v1.0 genome reference, the John Innes Centre Bioimaging facility and staff for their contribution to this publication, Matthew Hartley, John Innes Centre, is thanked for assistance in the analysis of starch granule-size distributions by coulter counter and the Biotechnology and Biological Sciences Research Council for funding via the Crop Improvement Research Club grant BB/J019496/1 and the Designing Future Wheat Institute Strategic Programme BB/P016855/1. DS thanks the Biotechnology and Biological Sciences Research Council (BBSRC, UK) for a Future Leader Fellowship BB/P010814/1.

## Supplementary material

Supplementary Table 1. Aegilops molecular markers.

Supplementary Table 2. Genes in the region containing *BGC1*.

Supplementary Table 3. PCR primers.

A. Cloning primers.

B. Primers used in Supplementary Figure 2.

Supplementary Table 4. Triple mutant genotyping primers.

Supplementary Figure 1. Aegilops *FLO6* cloning and analysis.

Supplementary Figure 2. Analysis of *FLO6* genes in *T. aestivum*, *Ae. peregrina* and KU37.

A. PCR analysis of *FLO6* genes.

B. The positions of the PCR products.

Supplementary Figure 3. *FLO6* TILLING lines of wheat.

A. The positions of FLO6 TILLING mutations.

B. Genotyping primers.

