## Supplementary Material for "A carbohydrate-binding protein, FLOURY ENDOSPERM 6 influences the initiation of A- and B-type starch granules in wheat"

Supplementary Table 1. Aegilops molecular markers.

Markers are designed for use in KASP assays except for 4G which was a COS marker used previously (Howard *et al*., 2011).

| Aegilops marker | Primer (VIC) | Primer (FAM) | Primer common | Wheat 4BS Ortholog |
|---|---|---|---|---|
| | VIC tag = GAAGGTCGGAGTCAACGGATT | FAM tag = GAAGGTGACCAAGTTCATGCT | | |
| TC60 | TCGTCGCCATGGAGGAGg | TCGTCGCCATGGAGGAGa | AGGACCAAAGACCGGGCG | TraesCS4B01G025500 |
| TC61 | ACAGTCTCCTAGGCGTCTGc | ACAGTCTCCTAGGCGTCTGa | CTACCAGCAGGAGAATAGGATC | TraesCS4B01G025600 |
| TC58 | AGTTGATACAGGTGCAGtTTa | AGTTGATACAGGTGCAGgTTc | CCATCTATTTGGCGGCAA | TraesCS4B01G026000 |
| KT108 | GGGAGATTGTGGTTATCTGGAAc | GGGAGATTGTGGTTATCTGGAAt | CAGCTGACTTCAATTCATTTAGC | TraesCS4B01G027900 |
| KT116 | GGAGATTGTGGTTATCTGGAAc | GGAGATTGTGGTTATCTGGAAt | CTGACTTCAATTCATTTAGCACTG | TraesCS4B01G027900 |
| KT113 | TCCGAgGTTCCAGAGCACGg | TCCGAgGTTCCAGAGCACGa | TGCTGTCAAGATGATTGTATGGAG | TraesCS4B01G028900 |
| TC36 | GCAGACTCAAACAACTTGCTc | GCAGACTCAAACAGCTTGCTt | CAGCTTTCTGAACTTGAGAGG | TraesCS4B01G029300 |
| TC35 | ATGTTGCCGTTGTAGTGGAc | ATGTTGCCGTTGTAGTGGAt | AGCACGCGGAGATCGACA | TraesCS4B01G029400 |
| KT117 | CTTCAAATAAATGGGGGCAc | CTTCAAATAAATGGGGGCAa | CCCAGTGGATGAGAATTTTC | TraesCS4B01G031000 |
| TC73 | GGATGGAACAATCAGAAAGc | GGATGGAACAATCAGAAGGa | ACCGGGATACAACCTCAGG | TraesCS4B01G029700 |
| KT70 | GCATGTCTTTAAGATATACATAAATaaatAAAC | GCATGTCTTTAAGATATACATAAATAAAC | AAGTAAGATGCCTTTCTGAAGTTCT | TraesCS4B01G030400 |
| KT110 | ATTTGGCATGCGGAATGGCTc | ATTTGGCATGCGGAATGGCTa | TTCATTCATACTTGATAATGCCC | TraesCS4B01G030500 |
| TC30 | GAAATCATTCGCCCCTGAc | GAAATCATTCGCCCCTGAa | GAGCTGCAGATTTGTTCCTG | TraesCS4B01G031300 |
| KT121 | TAGGCCCAGCACTGGTCAAc | TAGGCCCAGCACTGGTCAAa | TTACCTGAGATGTTTGATGACA | TraesCS4B01G032400 |
| TC34 | CAAGTACGGCCTCCCgAa | CAAGTACGGCCTCCCcAg | GGTGTAGGAGCTGACCGAG | TraesCS4B01G037000 |
| TC28 | GCCTTGCGCGCGAGgACc | GCCTTGCGCGCGAGcACg | AAGAACTCGGAGAAGCG | TraesCS4B01G033900 |
| TC26 | CTGGTGTACtATGGTCTGATCg | CTGGTGTACgATGGTCTGATCa | TAGCTTGTGTGGTTCATGTTAAT | TraesCS4B01G035600 |
| TC33 | GCATTTGAGGTGgAGGTCTg | GCATTTGAGGTGaAGGTCTa | CCAGAGAAGCAAGTGACCG | TraesCS4B01G037000 |
| 4G | GCAATCACGAACGGCTCGATCA | ATCTGGCAGCTTGCCAAGGCT | - | TraesCS4B01G037800 |

Supplementary Table 2. Genes in the region containing *BGC1*.

| Gene ID (Fig. 3) | Aegilops marker | 4A homoeolog | 4B homoeolog | 4D homoeolog | Annotation |
|---|---|---|---|---|---|
| 1 | KT113 | TraesCS4A01G284200 | TraesCS4B01G028900 | TraesCS4D01G026500 | Calcineurin-binding protein cabin-1 |
| 2 | | TraesCS4A01G282600 | TraesCS4B01G029100 | TraesCS4D01G028200 | F-box protein family-like |
| 3 | | TraesCS4A01G283500 | TraesCS4B01G029200 | TraesCS4D01G027100 | Glycine-rich family protein |
| 4 | TC36 | TraesCS4A01G283600 | TraesCS4B01G029300 | TraesCS4D01G027000 | Leucine-rich repeat protein kinase family protein |
| 5 | TC35 | TraesCS4A01G283700 | TraesCS4B01G029400 | TraesCS4D01G026900 | Hexosyltransferase |
| 6 | | TraesCS4A01G283900 | TraesCS4B01G029600 | TraesCS4D01G026800 | NRT1/PTR family protein 2.2 |
| 7 | TC73 | TraesCS4A01G284000 | TraesCS4B01G029700 | TraesCS4D01G026700 | Flo6 / 5'-AMP-activated protein kinase subunit beta-2 |
| 8 | | TraesCS4A01G283400 | TraesCS4B01G030100 | TraesCS4D01G027600 | Calcium-dependent protein kinase |
| 9 | | TraesCS4A01G258800 | TraesCS4B01G030200 | TraesCS4D01G027500 | Glycine-rich protein |
| 10 | | TraesCS4A01G283000 | TraesCS4B01G030300 | TraesCS4D01G027700 | B3 domain-containing protein |
| 11 | KT70 | TraesCS4A01G283200 | TraesCS4B01G030400 | TraesCS4D01G027800 | Inositol hexakisphosphate and diphosphoinositol-pentakisphosphate kinase |
| 12 | KT110 | TraesCS4A01G282800 | TraesCS4B01G030500 | TraesCS4D01G028000 | Tetratricopeptide repeat protein 38 |
| 13 | KT117 | TraesCS4A01G283100 TraesCS4A01G282900 | TraesCS4B01G031000, TraesCS4B01G035300LC | TraesCS4D01G027900 | Plasma membrane ATPase |

A region containing *BGC1* was defined by mapping in Aegilops (Fig. 1). Comparison of the Aegilops genetic map with the *T. aestivum* physical maps (Chinese Spring, Refseqv1) (IWGSC, 2018) identified 13 genes with orthologs in this region in all 3 genomes. The 13 genes were ordered according to their position in the wheat B genome and given ID numbers 1 to 13. Note that gene 13 occurs in duplicate on both 4AL and 4BS. The annotations of these gene are according to Ensembl Plants.

Supplementary Table 3. PCR primers.

A. Cloning primers.

The DNA polymerase used was Phusion HF (international.neb.com) and the PCR conditions were: (98 $^{o}$C for 30 s), 35 cycles (98 $^{o}$C for 10 s, 60 $^{o}$C for 30 s, 72 $^{o}$C for 15 s), (72 $^{o}$C for 10 min), 10 $^{o}$C hold. The clones were sequenced using the cloning primers and with additional internal primers (not shown).

| Forward primer | Sequence (5'-3') | Reverse primer | Sequence (5'-3') |
|---|---|---|---|
| KT106 KT Mf | CAGCAAGAATCAAGC | KT106 KT Mr | GCAGGATTGGGCCATACAAC |
| KT106 KT Ff | GCAGATGTGGTAAACGGGTT | KT106 KT Gr | TCCCATGGTTGTTCACGGTA |
| KT106 KT Hf | GCTCTGTGTTTGCCTGCTTA | KT106 KT Vr | CTTCTGCCGACCGATAGCTA |
| KT106 KT Wf | GAACAATCAGAAAGCGGCATTT | KT106 KT Wr | ACTAACCACGGACAACTTTGC |
| KT106 TC 4F | GGTGGCGTGCCTGTCAGT | KT106 TC 3R | AGATTACTCAAATAATTGCACTGCC |
| KT106 TC 5F | GAGCACATACAGGAATACAGGA | KT106 TC 19R | GCGATCTAATCAGTTGGCA |
| KT106 BS 1F | AGGTCGAGGAAGCGATCT | KT106 BS 2R | CGGCATGAGCTTAGCATTAA |

B. Primers used in Supplementary Figure 2A.

Template genomic DNAs were Chinese Spring (CS), KU37 (KU) and *Ae. peregrina* (AP). The predicted amplicon sizes are given. The DNA polymerase used was FastStart (sigmaaldrich.com) and the PCR conditions were: (95 $^{o}$C for 5 min), 45 cycles (95 $^{o}$C for 30 s; 60 $^{o}$C for 30 s; 72 $^{o}$C for 1.5 min), (72 $^{o}$C for 7 min), and 10 $^{o}$C to hold.

| PCR reaction | Designed to amplify | Primers (5' – 3') | Size | Template |
|---|---|---|---|---|
| **1** | Aegilops *FLO6-U* Intron 3 | GGCGCTCTATCGAAACATA + GCACGACGACAAAACACA | -<br>423 bp<br>439 bp | CS<br>KU<br>AP |
| **2** | Aegilops *FLO6-U* Exons 4 and 5 | TGGAAATATACATCGCTCTCG + GCTTGATGTTAGGGAGCATC | -<br>1396 bp<br>1397 bp | CS<br>KU<br>AP |
| **3** | Aegilops *FLO6-U* Exons 8 and 9 | ACAATTGAAATTAGTCAGCATAGG + CAGATGAGGAGTTCCTGGC | -<br>1045 bp<br>1050 bp | CS<br>KU<br>AP |
| **4** | Aegilops *FLO6-S* Intron 3 | CGGGCAGGAAGTGACGCT + TCCATCAATAACAAACCAAGG | -<br>802 bp<br>- | CS<br>KU<br>AP |
| **5** | Aegilops *FLO6-S* Exons 4 and 5 | GGAAATATACATCGCTCCCA + GCAGGCAGTGTAAGTTCATAGA | -<br>642 bp<br>- | CS<br>KU<br>AP |
| **6** | Aegilops *FLO6-S* Intron 5 | AATATTCCCTTGCAGTGCTC + TGCTTGATGTTAGGGAGAATG | -<br>479 bp<br>- | CS<br>KU<br>AP |
| **7** | Ta *FLO6 4D* and Aegilops *FLO6-S* Exons 8 and 9 | TCAGATCAGCCAGCAGAA + CCCGGGCTGGATCTTAGT | 1063 bp<br>949 bp<br>1080 bp | CS<br>KU<br>AP |

Supplementary Table 4. Triple mutant genotyping primers.

| Wheat line | Genome | Forward | Reverse | Amplicon size (bp) |
|---|---|---|---|---|
| Paragon deletion mutant. A-genome deletion | A | ATTGATGGACAACTAATGGATATT | AACACAGTGTTCAGGTCAACAT | 487 |
| Paragon deletion mutant D-genome deletion | D | GGGTTCTCTGATGATTGGG | AGCAGAGAATTGGGACATGG | 375 |
| Cadenza | B | GGGTTCTCTGATGATTGGG | GCCACCCATTTTGTAGTTAGTTAG | 423 |

Supplementary Figure 1. Aegilops *FLO6* cloning and analysis.

Alignment of predicted protein sequences. In rice *FLO6*, amino acids 1-71 encode the chloroplast transit peptide (predicted by ChloroP)

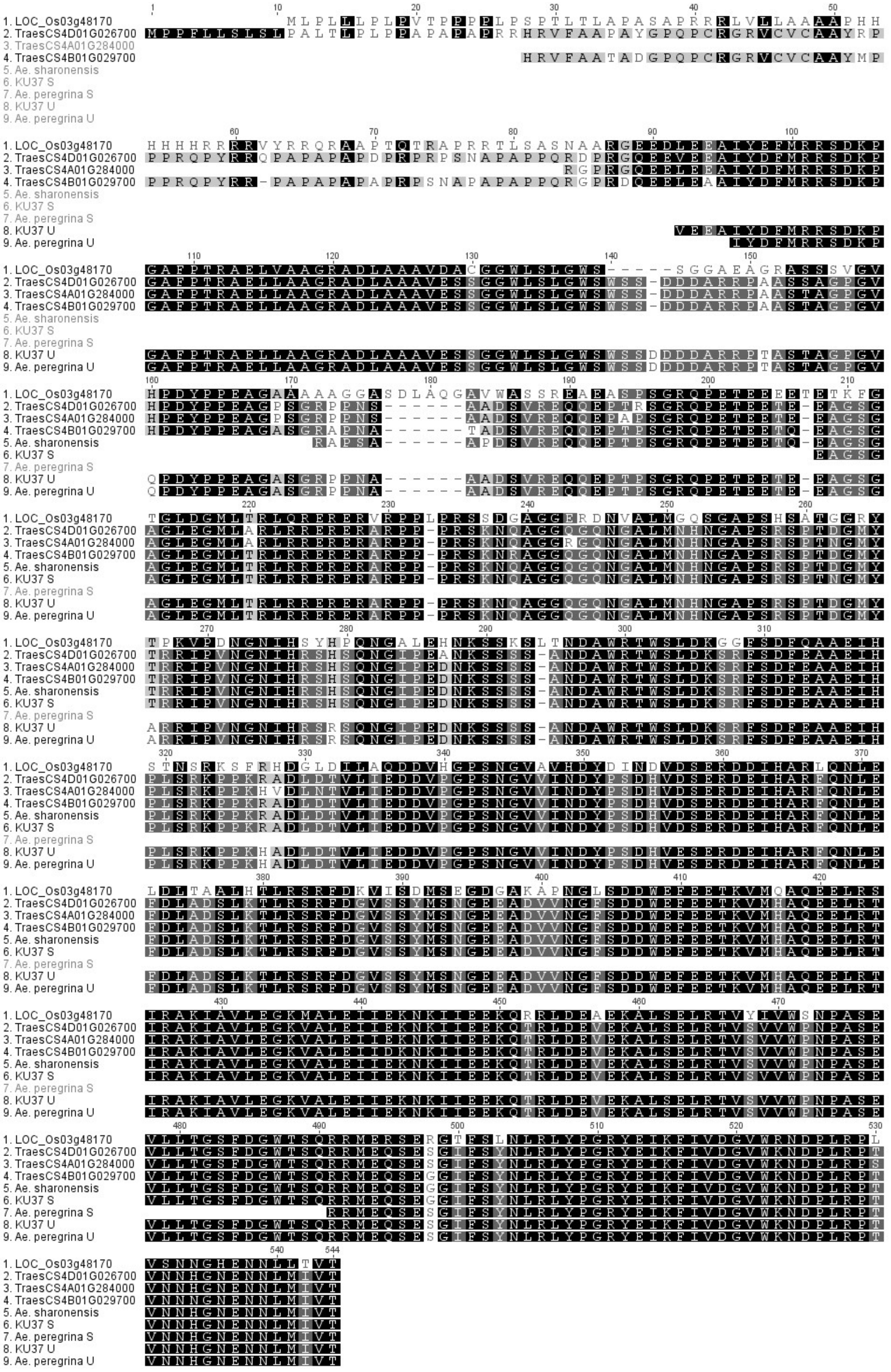

Supplementary Figure 2. Analysis of *FLO6* genes in *T. aestivum*, *Ae. peregrina* and KU37.

A. PCR analysis of the *FLO6* genes.

*FLO6* fragments were amplified by PCR using genomic DNA from wheat (Chinese Spring, CS), KU37 (synthetic tetraploid Aegilops with B-type granules, KU) and *Ae. peregrina* (natural mutant lacking B-type granules, AP). The primers were designed to amplify regions (PCR products P1-P7) of either *FLO6-U* (upper panel) or *FLO6-S* (lower panel) as indicated in B. Molecular weight markers are loaded at either side of each gel with sizes as indicated.

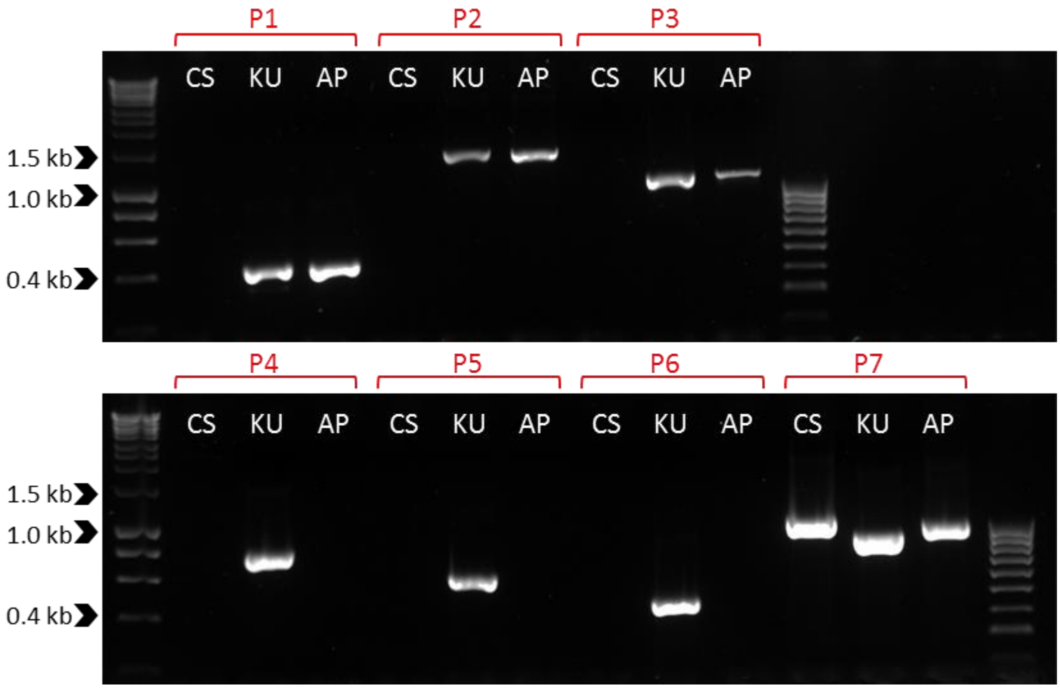


B. The positions of the PCR products in the *FLO6* genes.

The gene structure is as in Figure 4. P1-P7 are the PCR products shown in A.

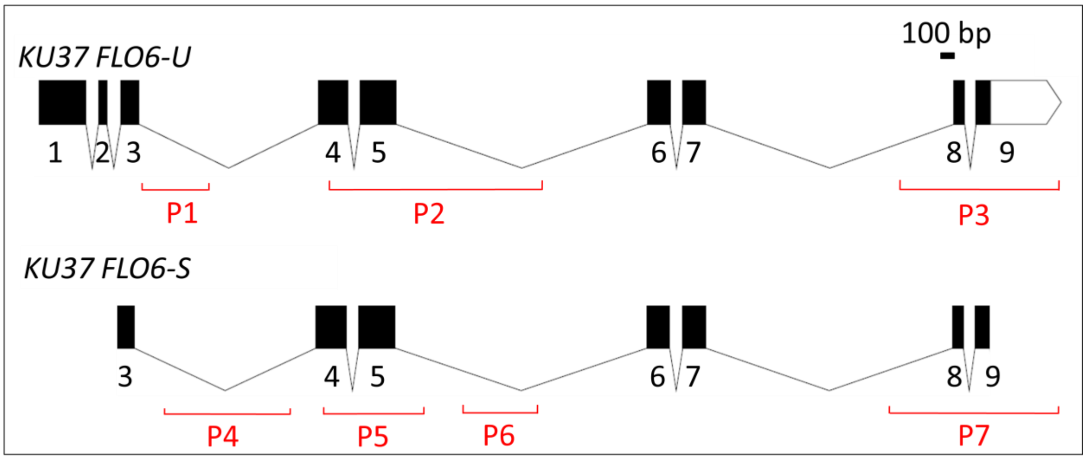

Supplementary Figure 3. The wheat *FLO6* TILLING mutants.

A. The positions of mutations in Kronos and Cadenza FLO6 genes (B) are indicated on the *FLO6-A* (*TraesCS4A02G284000*) and *FLO6-B (TraesCS4B02G029700*) gene structures. The effects of the mutations on the encoded proteins are indicated.

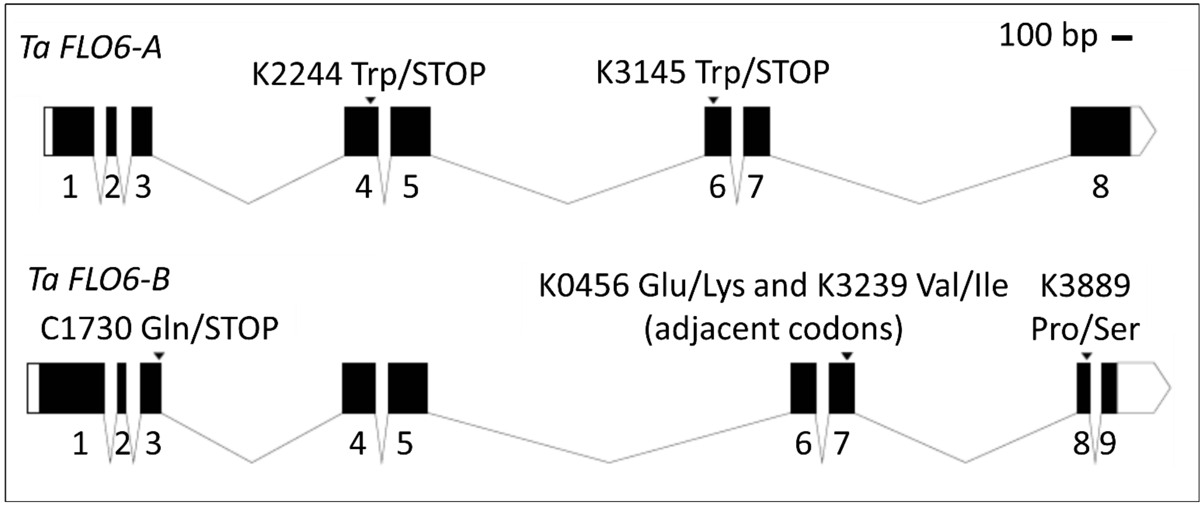


B. Genotyping primers.

| Cultivar (genome) | Chr | Mutant ID | Change to gene | Change to protein | Effect | Primer (VIC) | Primer (FAM) | Primer common |
|---|---|---|---|---|---|---|---|---|
| Kronos (AABB) | A | K2244 | G/A | Trp - STOP | nonsense | TTGTCAAGAGACCATGTTCGc | TTGTCAAGAGACCATGTTCGt | CTCCTAGTCGAAGTCCAACTA |
| Kronos (AABB) | A | K3145 | G/A | Trp - STOP | nonsense | TACTTTTGTCTCTTCAAATTCCc | ATTACTTTTGTCTCTTCAAATTCCt | GAAGATCTTGATGCATCTCATC |
| Kronos (AABB) | B | K0456 | G/A | Glu - Lys | missense | TATGGCCCAATCCTGCTTCAg | TATGGCCCAATCCTGCTTCAa | GCCACCCATTTTGTAGTTAGTTAG |
| Kronos (AABB) | B | K3239 | G/A | Val - Ile | missense | GCCCAATCCTGCTTCAGAAg | GCCCAATCCTGCTTCAGAAa | GCCACCCATTTTGTAGTTAGTTAG |
| Kronos (AABB) | B | K3889 | C/T | Pro - Ser | missense | CGCCATTACCTCATATCTACCAGg | CGCCATTACCTCATATCTACCAGa | ATAATCAAACACTGGACTTTTTCCT |
| Cadenza (AABBDD) | B | C1730 | C/T | Gln - STOP | nonsense | CGAGCGGGAGGGCAAGGTc | CGAGCGGGAGGGCAAGGTt | GCCGAGAGTTGAAAATGCACG |
